# From Stars to Molecules: AI Guided Device-Agnostic Super-Resolution Imaging

**DOI:** 10.1101/2025.02.25.640182

**Authors:** Dominik Vašinka, Filip Juráň, Jaromír Běhal, Miroslav Ježek

**Affiliations:** Department of Optics, Faculty of Science, Palacký University, 17. listopadu 12, 77900 Olomouc, Czechia

## Abstract

Super-resolution imaging has revolutionized the study of systems ranging from molecular structures to distant galaxies. However, existing super-resolution methods require extensive calibration and retraining for each imaging setup, limiting their practical deployment. We introduce a device-agnostic deep-learning framework for super-resolution imaging of point-like emitters that eliminates the need for calibration data or explicit knowledge of optical system parameters. Our model is trained on a diverse, numerically simulated dataset encompassing a broad range of imaging conditions, enabling generalization across different optical setups. Once trained, it reconstructs superresolved images directly from a single resolution-limited camera frame with superior accuracy and computational efficiency compared to state-of-the-art methods. We experimentally validate our approach using a custom microscopy setup with ground-truth emitter positions. We also demonstrate its versatility on astronomical and single-molecule localization microscopy datasets, achieving unprecedented resolution without prior information. Our findings establish a pathway toward universal, calibration-free super-resolution imaging, expanding its applicability across scientific disciplines.

## MAIN

Optical resolution is fundamentally constrained by diffraction. This limits the ability to observe structures comparable to the wavelength of light divided by the numerical aperture of the imaging system. Superresolution imaging has emerged as a transformative technique in disciplines ranging from biomedical [1, 2] and material sciences [3] to astronomy [4, 5], by circumventing this limit and revealing previously inaccessible structural details. Numerous applications rely specifically on imaging point-like or single-emitter sources. In biology, single-molecule localization microscopy enables nanoscale visualization of cellular structures [6–11]. Quantum physics leverages super-resolution for characterizing quantum dots [12] and precisely imaging cold atoms in optical lattices [13, 14]. Astronomy benefits through resolving individual stars and galaxies [15, 16].

Despite these successes, super-resolution methods require precise calibration and extensive knowledge of optical parameters. This requirement presents a substantial barrier, often involving laborious measurements, calibration data acquisition, and computationally expensive model retraining. Furthermore, practical applications frequently struggle with inhomogeneities and instabilities in imaging conditions, which further limit the versatility of existing algorithms.

To address these fundamental challenges, we propose a novel deep-learning framework that is inherently device-agnostic, eliminating the need for calibration or prior system-specific information. Our method exploits numerically simulated data encompassing an unprecedented range of optical conditions, enabling extreme generalization and adaptability across different imaging setups. Once trained, the model delivers rapid and accurate super-resolution reconstructions directly from a single, diffraction-limited image frame.

In this work, we demonstrate the efficacy of our approach through computational evaluations and experimental validations across multiple benchmarks, outperforming statistical Bayesian and deep-learning-based state-of-the-art methods. We further confirm the practical advantages and wide applicability of our approach using diverse datasets from molecular localization microscopy to astronomical imaging. The developed framework paves the way towards universal, calibration-free super-resolution imaging, significantly enhancing its accessibility and impact across scientific and technological disciplines.

### Super-resolution and device-dependence

Super-resolution imaging can be achieved through various approaches, such as linear inverse and statistical Bayesian algorithms [17], sparse representation [18], tomographic image synthesis [10, 19, 20], and methods based on blinking emitters [6– Many deep learning super-resolution methods have been developed in the last decade [21– Recently, artificial intelligence has been used to identify novel super-resolution microscopy setups [26].

Regardless of the specific approach, super-resolution imaging can generally be classified into two categories: reconstruction and parameter estimation. Reconstruction aims to directly restore the whole super-resolved image [17, 22], while estimation focuses on extracting key parameters and features, such as emitter localization [24, 27, 28]. The localization performance rapidly decreases for images with higher number of overlapping emitters, which typically limits its application to stochastically blinking or photoactivated samples. In the following discussion, we will focus exclusively on the former, more universal approach: image reconstruction.

Achieving accurate reconstruction of an object based on a single intensity image requires substantial knowledge about the imaging system under consideration. Central to this knowledge is an accurate estimation of the point spread function, which characterizes the response of the system to a point source of light. The reconstruction quality is inherently linked to the accuracy of the point spread function estimate; any inaccuracies will lead to poor reconstruction results [29]. However, its precise estimation entails additional calibration measurements involving an isolated emitting point source. This condition can prove very demanding, especially in systems with high concentration of emitters, low signal-to-noise ratio, or temporally developing systems. Additionally, imaging systems commonly exhibit variations in the shape of the point spread function across the image. Consequently, reconstructing larger areas may require separate calibration for each segment [30].

Some approaches can be adapted to perform a blind reconstruction, operating without explicit prior knowledge of the point spread function [31]. Instead, these methods impose additional assumptions on the imaging system, typically in the form of a predefined point spread function shape. During reconstruction, parameters of the point spread function, such as its width or other defining characteristics, are iteratively estimated directly from the input image. As a result, these methods introduce imperfections by simultaneously reconstructing the observed object and the point spread function of the imaging system. Altogether, blind reconstruction remains a non-convex optimization problem, posing a challenging task in real-life scenarios and often yielding inferior results compared to non-blind algorithms.

Similar device-dependent constraints arise with reconstruction using deep learning. While these approaches often outperform classical algorithms in both reconstruction and parameter estimation tasks [21, 22, 24, 25], their calibration presents an even more significant obstacle. Unlike traditional approaches, deep learning model operates on a data-driven paradigm, inferring the reconstruction mapping by extracting relevant information from observed samples. However, due to the device-dependent nature of a typical training process, the model can only be applied to a particular point spread function profile and additional conditions such as emitter power, background noise distribution, and concentration of emitters, each requiring prior estimation from the calibration datasets [32]. Modifying these parameters requires remeasuring the calibration set and retraining the model, making it a resource-expensive procedure.

## RESULTS

### Device-agnostic modeling

We developed a device-agnostic modeling network (DAMN) designed to reconstruct an intensity image of point-like emitting sources, as illustrated in Fig. 1. Unlike traditional approaches, the DAMN model operates using a single intensity frame without requiring additional information about the optical parameters of the imaging system. Without the need for calibration measurements or retraining, it can directly process images from diverse applications. Our model leverages the convolutional architecture of a neural network (refer to the Methods section), enabling it to reconstruct frames of varying shapes and sizes. Moreover, the reconstruction process is performed by a single-pass propagation through the neural network, resulting in rapid reconstruction of super-resolved images.

**FIG. 1.**
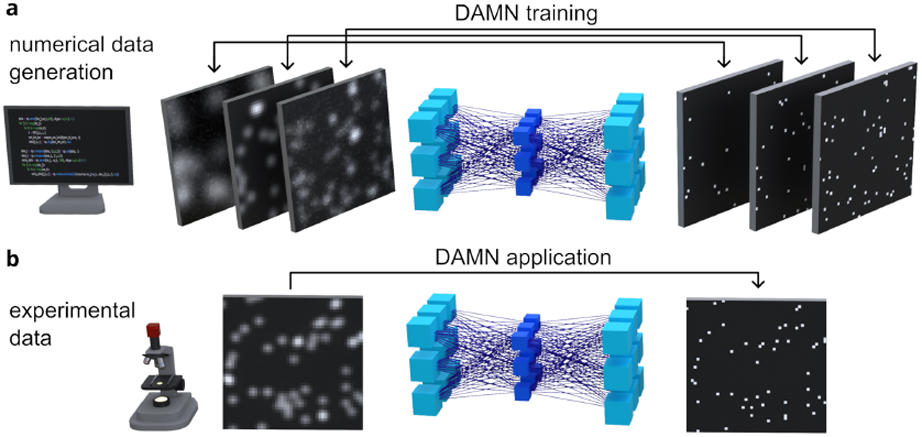
Schematic representation of the DAMN approach and its application to intensity images of point-like emitting sources. **a** The model is trained using numerically simulated data pairs comprising resolution-limited noisy images alongside their super-resolved counterparts. Each training sample represents a unique combination of underlying optical parameters, such as the width of the point spread function and the signal-to-noise ratio. **b** Following training, this model is applied to enhance the resolution of experimental images acquired using a real-life imaging system.

We train the DAMN model using pairs of targeted ideal objects and their corresponding resolution-limited images generated through numerical simulation. The ensemble of data pairs covers various optical parameters, such as emitter power and concentration, background noise distribution, and even different shapes and widths of the point spread function. Their ranges were selected to cover the majority of realistic situations (additional details in the Methods section). The extent of the synthetic training data is unprecedented in the astronomy or emitter microscopy fields. Subsequently, we utilized incremental learning techniques [33] for the model to acquire knowledge about the widest possible parameter combinations. As a result of the optimization and training process, this single model accurately reconstructs superresolved images independently of the underlying parameter values. Consequently, without the need for retraining or calibration data, the model can be applied to diverse imaging systems and is resistant to their parameter changes, including inhomogeneity across a field of view and instability, such as drifts and fluctuations in time.

### Evaluation using simulated datasets

To characterize the effectiveness of the DAMN model, we conduct a comparative analysis with the Richardson-Lucy deconvolution based on Bayesian estimation with a uniform prior [34, 35]. This widely used iterative algorithm implements image reconstruction by repeatedly utilizing a known point spread function. We evaluate the results using a separate test set of image pairs not included in the training data to assess the generalization ability for new samples. We quantify the performance of both methods using the mean absolute error, calculated between the reconstructed image and its target object, averaged over the whole test dataset. In contrast to the Richardson-Lucy algorithm, which requires (and was supplied with) prior knowledge of the imaging system, our DAMN model operates independently of any device-dependent information. Despite this distinction, the device-agnostic model (without the prior information) outperforms the classical Richardson-Lucy deconvolution (with complete information on the point spread function) in terms of reconstruction quality, as discussed in the following text. Additionally, due to the iterative nature of the Richardson-Lucy algorithm, it requires three orders of magnitude higher computational time on average, further emphasize the advantages of the DAMN approach, see the Methods section.

Panels **a c** in Fig. 2 illustrate the dependence of mean absolute error on the emitter power, the width of the point spread function, and the concentration of emitters, respectively. The results of these log-log graphs are characterized using an average value over the test dataset with a corresponding 90% confidence interval. Panel **a** contains dual horizontal axes representing the varying emitter power using both the signal-to-noise ratio (SNR) and the peak-to-noise ratio (PNR). For additional details regarding the explored parameters and evaluation of both approaches, see the Methods section. The DAMN model, represented by the red color, exhibits significantly better performance than the green-colored Richardson-Lucy algorithm. Similar results can also be observed in panels **b** and **c**. Panel **b** depicts the error dependence on the point spread function width (consult the Methods section). The remaining panel **c** characterizes the dependence on the emitter concentration, referring to the collective number of emitters in a single image. As seen across all three panels, our device-agnostic approach provides up to two orders of magnitude more accurate reconstructions than the device-dependent Richardson-Lucy deconvolution. The depicted errors were evaluated across the whole 50 *×* 50 pixel image and can be, alternatively, expressed as per-pixel errors by dividing the values by the number of pixels.

**FIG. 2.**
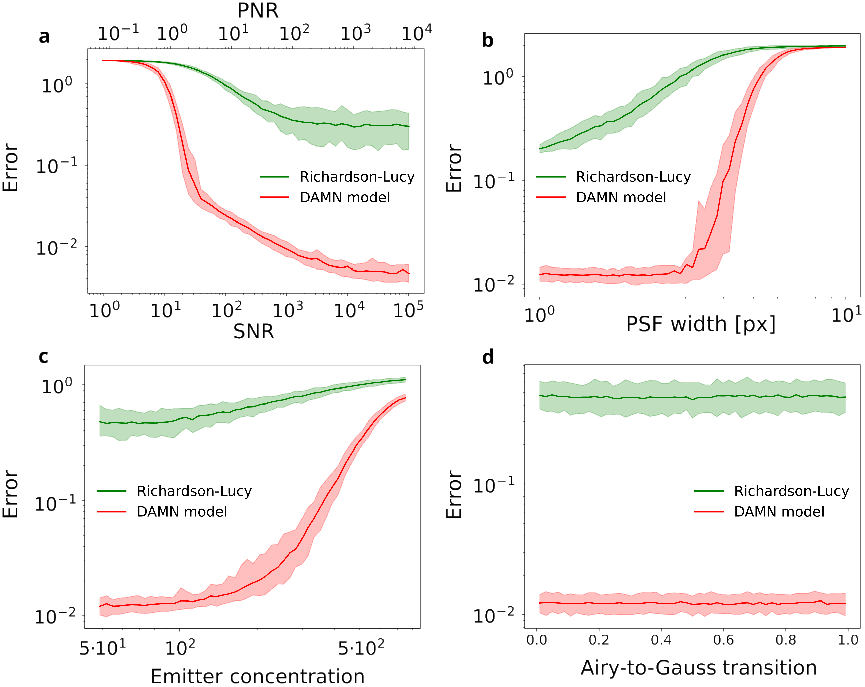
The dependence of the mean absolute error on **a** the signal-to-noise ratio (SNR), **b** the width of a Gaussian point spread function (PSF), **c** the number of emitters in the image (concentration), and **d** the continuous transition between an Airy and a Gaussian PSF, respectively. The resulting averages of the Richardson-Lucy algorithm (green) and the DAMN model (red) are accompanied by their 90% confidence intervals over the test set. Panel **a** is provided with a secondary horizontal axis recalculating the SNR values to the peak-to-noise ratio (PNR). Across all panels, the DAMN model consistently outperforms the Richardson-Lucy deconvolution by up to two orders of magnitude. The optical parameters not investigated in a given panel have the following values: SNR = 500, the average noise intensity = 10, the concentration = 50, and the Gaussian PSF with *σ* = 2 px.

The panel **d** in Fig. 2 visualizes the error dependence on the point spread function shape, gradually transitioning from an Airy pattern to a Gaussian profile of the same width. As seen from this semi-logarithmic graph, the performance of both methods remains approximately constant during this transition. Such behavior is expected from the Richardson-Lucy deconvolution, as we always provide the correct point spread function profile. On the other hand, the DAMN model was trained using solely the exact Gaussian and Airy point spread functions constituting half of the dataset each. Despite this simplification in the training process, the graph unambiguously demonstrates the adaptive ability of the DAMN approach to generalize to previously unseen point spread function shapes. This adaptability is beneficial during training, as it significantly reduces the data and simulation complexity required for training device-agnostic models. Altogether, the presented panels display numerous benefits of the DAMN approach, proving it superior to the stateof-the-art algorithm.

### Optical microscopy experimental validation

Demonstrating the validity of our approach beyond the simulated data, we applied it to reconstruct experimentally acquired images. Explicitly for this, we developed a custom-built optical microscopy setup illustrated in Fig. 3 and shown in detail in Extended data Fig. 1.

**FIG. 3.**
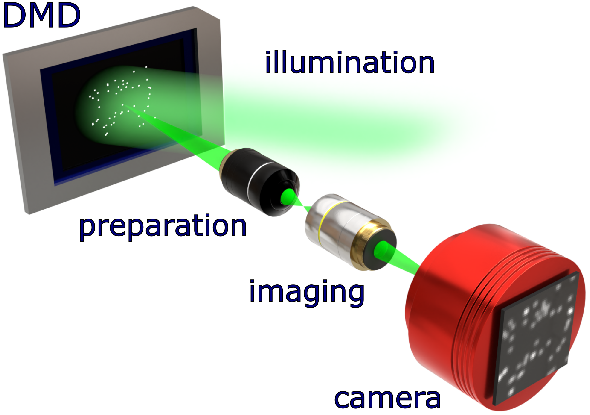
Schematic illustration of the optical setup used to collect experimental data pairs. A ground-truth mask is imposed on the digital micromirror device (DMD) by configuring its mirrors. An incoherent illumination light reflected by these mirrors impinges a high-resolution preparation system. The DMD-imposed mask is re-imaged into the front sample plane of the preparation system, creating point-like emitters with the intended spatial distribution. The imaging part of the setup, comprised of a low-resolution microscope objective, images the sample-plane emitters onto a camera. The resulting camera-captured intensity image and the DMD-imposed mask represent the experimental data pairs.

This microscope provides complete control over the spatial distribution of emitting point-like sources, achieving this for the first time in super-resolution benchmarking. An incoherent light illuminates a digital micromirror device (DMD), onto which a mask representing the targeted ground-truth image is imposed. By configuring the micromirrors, we prepare a mask containing an arbitrary number of sources positioned at desired locations. This mask is then re-imaged by a high-demagnification and high-resolution preparation system, creating point-like emitters and thus forming a sample object. Subsequently, a low-resolution imaging microscope projects the emitters onto a camera. The resulting intensity profile captured by the camera serves as the resolution-limited input for reconstruction methods. Together with the groundtruth mask, this setup provides experimental test data pairs, which allow for exact metric quantification of each reconstruction method performance. For details, see the Methods section.

Using this setup, we investigated the performance of both methods while varying the concentration of emitters. Panel **a** in Fig. 4 illustrates the mean absolute error between reconstructed images and their corresponding target objects using the experimental data, shown as dot markers. For comparison, the continuous lines represent the results obtained using simulated data with corresponding optical parameters. The DAMN model outperforms the Richardson-Lucy deconvolution by orders of magnitude, even when applied to real-world images acquired from experimental measurements. This improvement is achieved despite operating significantly below the resolution limit of our optical system. To further emphasize these findings, panel **b** in Fig. 4 shows a representative experimental image containing nearly 200 emitters and its reconstruction by each method. As observed, the DAMN model offers a near-perfect reconstruction of the original image without the need for any calibration. The Richardson-Lucy deconvolution exhibits severe deviations and artifacts despite knowing the point spread function of the imaging setup. This experimental demonstration highlights the benefits of our device-agnostic approach, which significantly outperforms the established state-of-the-art algorithm requiring prior knowledge of the point spread function.

**FIG. 4.**
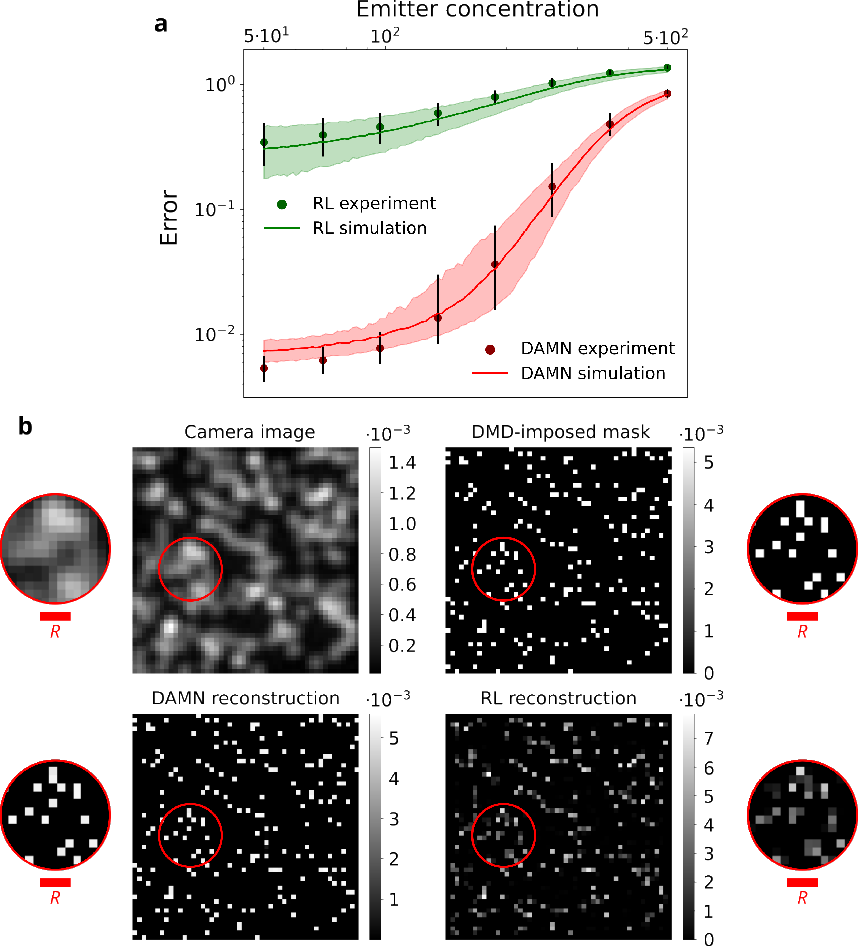
**a** The red and green dots represent the mean absolute error between the DMD-imposed masks and the superresolved images reconstructed by each method. These errors were evaluated across various emitter concentrations. The accompanying continuous lines depict error values derived from simulated data using optical parameters estimated for our imaging system. **b** A typical camera image containing nearly 200 emitters, alongside its corresponding DMD-imposed mask and each method reconstructions. The circled areas contain a magnified region for easier visual comparison. It is evident that the DAMN model significantly outperforms the Richardson-Lucy (RL) algorithm even in regions where mutual emitter distance is well-below the Rayleigh resolution limit *R* = 3.9 px (inset scalebars).

### Astronomical and localization microscopy demonstration

We have also explored the benefits of incorporating upsampling layers into the DAMN convolutional architecture, further extending its capabilities. The modified network reconstructs images with an eightfold increase in dimensions, leading to an even more prominent improvement in the resolution. We applied this model to an astronomy image of a dense star cluster from observations of the southern spiral galaxy NGC 300 in the Sculptor constellation by the European Southern Observatory [36], see Fig. 5 **a**. Panel **b** depicts a magnified segment of the galaxy with marked regions of interest, while panel **c** presents their high-resolution reconstruction produced by the DAMN model. Despite the stars in all three regions being below the resolution limit of the imaging system, the DAMN reconstruction clearly reveals them. Notably, a detailed examination of region III shows two distinct intensity peaks. While the original data resemble a single structure, our reconstruction indicates that it might be two proximal stars, with an approximate 0.11 arcsec angular distance, hidden deep below the resolution limit. Confirming this hypothesis would require further analysis using additional data. While the DAMN model can process the entire 7600 × 7600 pixel image, saturation artifacts in the central part of the original data (Fig. 5 **a**) led us to demonstrate the reconstruction of smaller outer regions of the galaxy. Moreover, as shown in **c**, the resolution enhancement is so significant that proper visualization requires considerable zoom into the reconstructed areas.

**FIG. 5.**
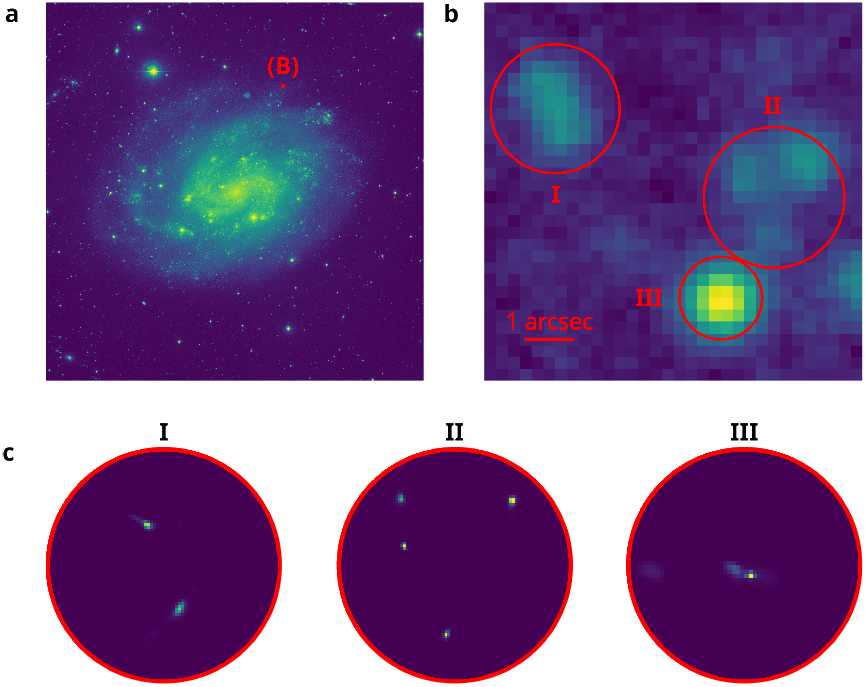
Demonstration of the DAMN model super-resolving capabilities on an astronomy image of a dense star cluster. **a** Intensity image of the spiral galaxy NGC 300 acquired by European Southern Observatory. **b** Magnified region of the galaxy, highlighting key areas of interest. **c** High-resolution reconstructions of these regions generated by the DAMN model, revealing details of individual stars. These reconstructions do not use any calibration data or prior information on the employed imaging system. The region III reconstruction indicates the presence of a secondary source completely beyond the optical resolution of the telescope.

Additionally, we applied the same DAMN model, without retraining, to a publicly available dataset [37, 38] acquired for the single-molecule localization microscopy challenge. Fig. 6 depicts the performance evaluation using these 500 tubulin images of 128 × 128 pixels with high molecule concentration **a**. Panel **b** shows a reference image provided by the SOSplugin [39], a least-square localization method assuming a Gaussian point spread function, which participated in the challenge and is available as a plugin for ImageJ. Panel **c** depicts a tubulin reconstruction from Deep-STORM [22], a stateof-the-art deep learning model for image reconstruction in localization microscopy. Unlike our approach, DeepSTORM operates in a device-dependent regime, requiring extensive optical system information, such as camera specification, point-spread-function model, approximate SNR, and expected emitter density. In comparison, the DAMN model directly processes the dataset without calibration or prior knowledge of the imaging system and provides the super-resolved tubulin image, see panel **d**. The highlighted areas show a close-up comparison of all three approaches. Our DAMN model accurately captures the underlying tubulin structure, achieving resolution significantly surpassing that of the SOSplugin localization method. Unlike the localization method, the DAMN model clearly resolves the two vertically oriented, overlapping microtubules and reconstructs the horizontal one without gaps. Moreover, a close inspection reveals that our model provides a sharper image with a cleaner and thinner microtubule structure compared to the device-dependent Deep-STORM. Panel **e** visualizes the projection of the microtubule profile over the yellow segment, demonstrating the resolution improvement. This result ultimately underlines the reconstruction ability of device-agnostic models in real-world applications. Although the presented model provides super-resolved reconstructions, the same paradigm of device-agnostic deep learning can be applied to localization tasks, i.e., directly predicting the positions and intensities of the emitting sources. Such a model could be highly advantageous for single-molecule localization microscopy of densely populated samples.

**FIG. 6.**
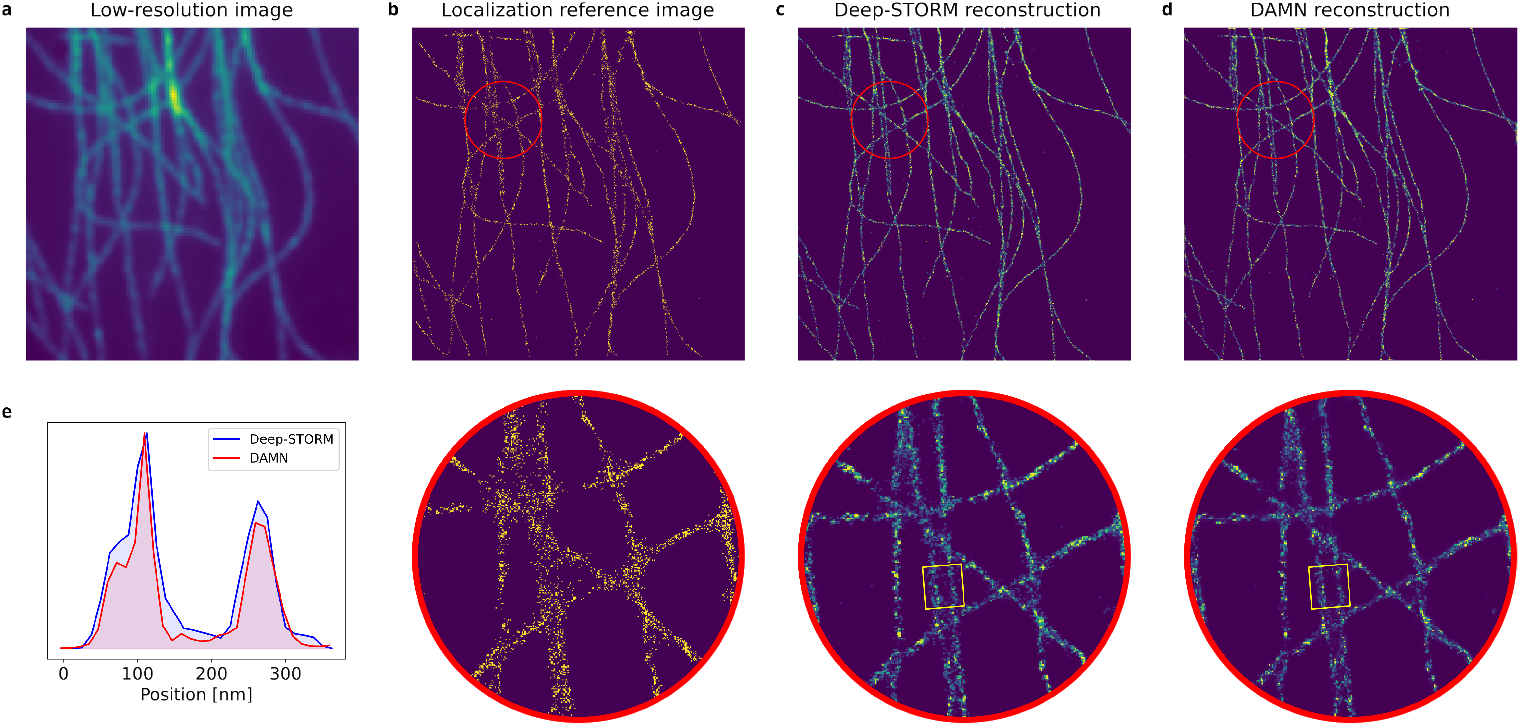
Demonstration of the DAMN model on a high-concentration tubulin dataset from the single-molecule localization microscopy challenge. **a** A low-resolution image integrated from the set of 500 measured images, containing numerous emitting molecules. **b** A reference binary map of localized molecule positions derived from the dataset. **c** A reconstruction generated by Deep-STORM, a device-dependent deep learning approach calibrated using detailed information on the imaging setup. **d** A super-resolved image produced from the calibration-free, device-agnostic model without prior information. Panels **b-d** include inset images with a magnified region highlighting the resolution details. **e** Projection of the microtubule profile over the yellow rectangle segment.

## DISCUSSION

We demonstrated a device-agnostic approach to superresolution imaging of point-like emitting sources that utilizes deep learning techniques. The presented approach solves the long-standing problems of measuring calibration data and estimating parameters accompanying the training of a deep learning model. Using only numerically simulated data, we develop a model capable of reconstructing intensity images using a single frame without explicit knowledge of the imaging system. This model can be applied to images of arbitrary shape and size originating from various optical systems, and the calibration-free nature grants it resistance against non-uniformity and drifts of the optical parameters. We compared this device-agnostic model with the well-established Richardson-Lucy deconvolution algorithm. The analysis results undeniably show the superior performance of our approach, outperforming the Richardson-Lucy algorithm by orders of magnitude in terms of both reconstruction accuracy and speed. Moreover, we designed an advanced optical setup for acquiring experimental images of emitting sources together with their precise ground-truth references. Providing full control over the spatial distribution of emitters, this setup enables exact performance quantification of any reconstruction method. Using these experimental data pairs, we verified the superior performance of our approach.

To further demonstrate the universality of our approach, we applied the model to astronomy and microscopy data, achieving significant resolution improvements in both domains. We reconstructed a highresolution tubulin image from single-molecule localization microscopy data. While in astronomy, we enhanced the resolution of a densely packed star cluster image. Notably, all DAMN reconstructions were performed without any prior knowledge of the optical setup, data preprocessing, or parameter estimation. Despite this, the obtained results surpass the state-of-the-art methods. With sufficient computation resources, even further improvements can be achieved by incorporating more complex data simulations and larger network architectures. By taking this first step, our work lays a foundation for universal image reconstruction, entirely independent of optical settings, thus contributing to the advancement of image processing in various fields.

## METHODS

### Data Simulation

A resolution-limited image can be characterized by its underlying optical parameters. In our case, these parameters include the shape and width of the point spread function, the background noise intensity, the emitter power representing the number of emitted photons, and the number of emitters present within a 50 × 50 pixels field of view of an image, which we term emitter concentration. Consequently, all possible resolution-limited images form a high-dimensional space. To implement the device-agnostic framework with a data-driven deep learning algorithm, the model must observe data samples covering the space during training. Therefore, the approach highly benefits from using simulated data pairs, as collecting a sufficient amount of experimentally acquired samples would be exceptionally time-consuming. Additionally, data simulation allows using incremental learning techniques (see Deep Learning Model), which are ideal for applications with large datasets. The following ranges of optical parameters were used for the data simulation: the emitter power ∈ [1, 10^5]^, the average noise ∈ [1, 100], the emitter concentration ∈ [5, 500], and the point spread function width ∈ [10^*−*0.25^, 10^1.25^] px ≈ [0.5, 17.75] px. For the first time in the emitter visualization community, data with such a broad scale of imaging parameters have been generated and used to train an artificial neural network.

The generation process of a single simulated data sample follows these steps. First, we generate the emitter concentration, followed by assigning each emitter its power and pixel position in the image. Next, we perform a convolution with the point spread function to simulate the effects of finite resolution. Subsequently, shot noise is added to each pixel of the image. With knowledge of the average noise intensity and emitter power, we can calculate the signal-to-noise ratio of the generated sample as 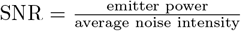. Alternatively, we can calculate the peak-to-noise ratio, 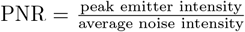, using the maximum value of the convolved emitter intensity. It is worth noting that the simulation process can be further enhanced by incorporating additional features and parameters, such as point spread function asymmetry and aberrations [40, 41]. For this showcase study, we opted to simplify the simulation to reduce the computational demand during training. Despite this simplification, the DAMN model achieves excellent results on experimental data and demonstrates significant generalization ability beyond the expected regime, as shown in Fig. 2 **d**.

To perform the convolution, we generate the convolution matrix of the point spread function using two distinct parameters shape and width. The shape parameter distinguishes between an Airy pattern *A*, simulating low numerical aperture scenarios, and a two-dimensional Gaussian pattern *G*, typically used for cases with higher numerical aperture values [42]. The convolution matrix is normalized to a unit sum to conserve the emitter power. The pattern functions can be expressed as

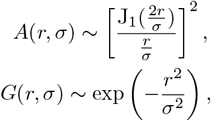

where *r* is the distance from the center, and J_1_ is the first-order Bessel function of the first kind. The variable *σ* is the width parameter, which dictates the full width at half maximum of the point spread function as 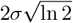.

### Deep Learning Model

The DAMN model is a deep convolutional neural network comprising 35 hidden layers with 71 channels each and a final output layer with a single channel. Information flows through each hidden layer via trainable 7 × 7 filters that apply local convolutional operations, allowing the network to detect localized patterns. These transformations are followed by a LeakyReLU activation function with a 0.05 negative slope, which introduces a computationally efficient non-linearity without causing the dying ReLU problem [43]. Since we normalize the input samples to a unit sum before processing, the output layer utilizes a softmax activation function [44] to preserve this condition and ensure non-negative values. To prevent overfitting, each hidden layer is paired with dropout regularization at a 0.01 rate, randomly setting a fraction of input units to zero during training [45]. This regularization encourages the network to learn robust features that do not rely on any specific neurons. As the DAMN model consists solely of convolutional layers, it can process images of arbitrary shapes and sizes without retraining. Additional resolution improvement is achieved by incorporating upsampling layers into the convolutional architecture. These layers increase the image dimension by repeating its rows and columns. The modified model applied in Fig. 5 and 6 contains three upsampling layers that double the input dimensions, resulting in their eightfold increase. This architecture makes the network highly flexible and effective for image-processing applications.

The presented model architecture results from extensive optimization utilizing a mesh adaptive direct search using the Nomad library [46], as well as numerous additional manual adjustments. Altogether, our model contains over 8 million trainable parameters and was trained incrementally on nearly 3 million simulated images, with a 25% validation set split. Incremental learning techniques gradually present the training data to the network. These techniques involve training the model using randomly generated data, which are continuously replaced with newly generated samples throughout the training process [33]. The level of network complexity and dataset size is unprecedented in the field of emitter reconstruction and localization. Nevertheless, the model remains relatively modest compared to modern large language models, suggesting further potential to integrate additional model features by increasing complexity.

The training of the DAMN model follows the backpropagation algorithm, which computes the gradient of the loss function with respect to each weight in the network using the chain rule. We use a mean squared error (MSE) loss, calculated between the predicted *I*_1_ and target *I*_2_ images and averaged over a batch of data samples. Given a set of two-dimensional image pairs, we calculate the loss as

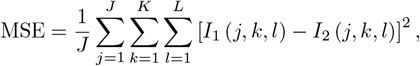

where *J* is the batch size, *K* and *L* are the image dimensions, and *I*(*j, k, l*) is the intensity of the *j*-th image in the batch at the pixel position (*k, l*). The gradients obtained from this loss are used to iteratively update the weights with each batch of 128 training samples. To improve convergence towards the minimum loss, we implement the Adam optimizer [47], which incorporates adaptive moment estimation. Adam updates the weights using both the first and second moments of the gradient, making the descent more efficient. A portion of the training data is reserved as a validation set to actively monitor the convergence using the mean absolute error (MAE) metric

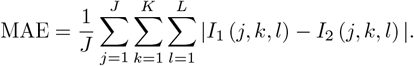

The metric provides an additional measure of model performance that is, unlike a loss function, not directly optimized during training.

### Richardson-Lucy Algorithm

The Richardson-Lucy algorithm is an iterative method used to restore an image blurred by a known point spread function [34, 35]. This sophisticated and highly flexible algorithm is derived from the Bayesian probability theory with a flat prior. It maximizes the posterior probability that the observed image results from the estimated image convolved with the given point spread function. Assuming a multinomial distribution of detection events, the Richardson-Lucy is equivalent to the expectation-maximization algorithm for likelihood maximization in positive linear inverse problems [48, 49]. The quality of the reconstructed image relies heavily on accurate knowledge of the point spread function; discrepancies can lead to artifacts and inaccurate reconstructions. Since we simulate the testing data, the Richardson-Lucy algorithm can use the precise point spread function, making it an excellent method for providing a state-of-the-art baseline for evaluating the DAMN model.

In addition to the point spread function *P*, the iterative algorithm requires an initial guess for the reconstruction, *I*^(0)^. A common practice is to set this guess as the observed resolution-limited image *I*_blurred_ or to start with a uniform image. We explored both approaches and chose to set *I*^(0)^ = *I*_blurred_, as there was a negligible difference in accuracy and computational speed. Then, an iterative procedure updates the estimation at the *k* + 1 step

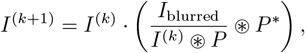

where ⊛ is convolution, and *P* ^*\**^ denotes the flipped point spread function. Both the initial guess *I*^(0)^ and the point spread function *P* need to be normalized to a unit sum. We implement two stopping criteria for the algorithm: either the mean absolute error between successive iterations becomes smaller than 10^*−*10^ per pixel, or the number of iterations exceeds 10^6^. The first criterion is typically met in the majority of cases.

Furthermore, we evaluated the computation time required by the Richardson-Lucy algorithm. As the time depends on the underlying optical parameters of an image, we have simulated an extensive test dataset containing approximately 300,000 randomly generated samples. Using an AMD EPYC 7443P 24-core processor, the Richardson-Lucy needs almost five days to reconstruct this dataset. In comparison, the DAMN model utilizing the same CPU resources reconstructs the data in a little over two hours. Moreover, the DAMN model with a GPU-enabled operation requires only approximately eight minutes for the same task when using an NVIDIA RTX A5500 GPU.

### Experiment

The optical setup, which provides resolution-limited experimental images with their corresponding target objects, consists of three segments, see Extended Data Fig. 1. The first segment prepares the illumination of the digital micromirror device (DMD). An incoherent broadband light with a central wavelength of 555 nm, produced by a laser-induced fluorescence light source (Crytur Mon-aLIGHT B01), is coupled to a 4 mm diameter light guide. The emerging light is collimated by a microscope objective MO_s_ (OLYMPUS 20x/0.4). An iris diaphragm D is positioned proximal to the objective. This diaphragm is re-imaged onto the DMD (Texas Instruments DLP LightCrafter 6500) by a telescope consisting of two achromatic doublets, L_s1_ and L_s2_, with focal lengths of 50 mm and 150 mm, respectively. The telescope improves the illumination homogeneity, and re-imaging the diaphragm reduces reflections from the passive parts of the DMD.

In the second setup segment, point-like emitters with the targeted spatial distribution are prepared. By addressing the DMD, we impose a ground-truth mask with each pixel comprising a 5 × 5 micromirror grid with a 7.6 *µ*m micromirror pitch. As we address only the central micromirror of the grid, the closest distance between two emitting sources is 38 *µ*m in the DMD plane, see Extended Data Fig. 1. This mask is re-imaged into the front focal plane of MO_p_ by a preparation telescope consisting of a 200 mm focal-length achromatic doublet L_p_ and a high-resolution microscope objective MO_p_ (OLYMPUS 100x/0.9). As the point-like emitters, of the approximate 310 nm full width at half maximum and 340 nm minimal distance, are created in this plane, we term it the emitter plane.

Unlike the first two segments, which prepare the emitters and provide complete control over their spatial distributions, the final segment represents an imaging setup with optical parameters unknown to the DAMN model. A low-resolution microscope objective MO_I_ (OLYMPUS 10x/0.25) projects the emitter plane onto a CMOS camera (ZWO-ASI 178MM, 2.4 *µ*m square pixels) with a mounted spectral filter (central wavelength of 532 nm with 10 nm spectral width). The MO_I_ resolution is approximately 1.13 *µ*m, regarding the full width at half maximum in the emitter plane. In comparison, the 340 nm minimal emitter distance is 3.3× below this resolution limit. The emitter plane is imaged with an approximate 230 × magnification to match the major part of the camera chip. The exposure time for each measurement is set to 5.7 seconds to utilize the majority of the 14-bit dynamic range of the camera while avoiding its saturation. The captured image is down-scaled by a factor of 32 to match the 50 × 50 ground-truth mask size to enable metric evaluations. Using this imaging setup, we collected the experimental dataset of resolution-limited camera images and their corresponding ground-truth masks for various emitter concentrations, see Fig. 4 **a**. Additionally, we measured a high-SNR calibration set for precise estimation of the point spread function, which we provided only to the Richardson-Lucy algorithm. Besides the concentration ranging from 50 to 500 emitters, the optical parameters in the collected dataset exhibit the following estimated values: SNR = 2300, average noise intensity = 10, and an Airy-shaped kernel with *σ* = 2.05 px, equivalent to the 3.4 px full width at half maximum. The Rayleigh resolution limit of our imaging system, given by the radius R of the first dark ring in the Airy pattern, is approximately 3.9 px, see Fig. 4 **b**.

## DATA AVAILABILITY

The code and data supporting the results of this study are publicly available on GitHub [50]. University Olomouc. We thank J. Provazník for maintaining the cluster and providing support.

## AUTHOR CONTRIBUTIONS

M.J. conceived the idea of a device-agnostic approach to image super-resolution and supervised the project. D.V. developed deep-learning models and performed numerical simulations and data processing. F.J. and J.B. developed the optical experiment, performed data acquisition, and participated in data processing. J.B. supervised the experimental part of the project. D.V. wrote the manuscript, and all authors were involved in revising the manuscript.

## FUNDING

This work was supported by the Czech Science Foundation (project 21-18545S), and the Ministry of Education, Youth, and Sports of the Czech Republic (project OP JAC CZ.02.01.01/00/23 021/0008790). DV acknowledges the support by Palacký University (projects IGA-PrF-2024-008 and IGA-PrF-2025-010).

## DISCLOSURES

The authors declare no conflict of interest.

## ACKNOWLEDGMENTS

We acknowledge the massive use of cluster computing resources provided by the Department of Optics, Palacký

---

**Extended Data Fig. 1.**
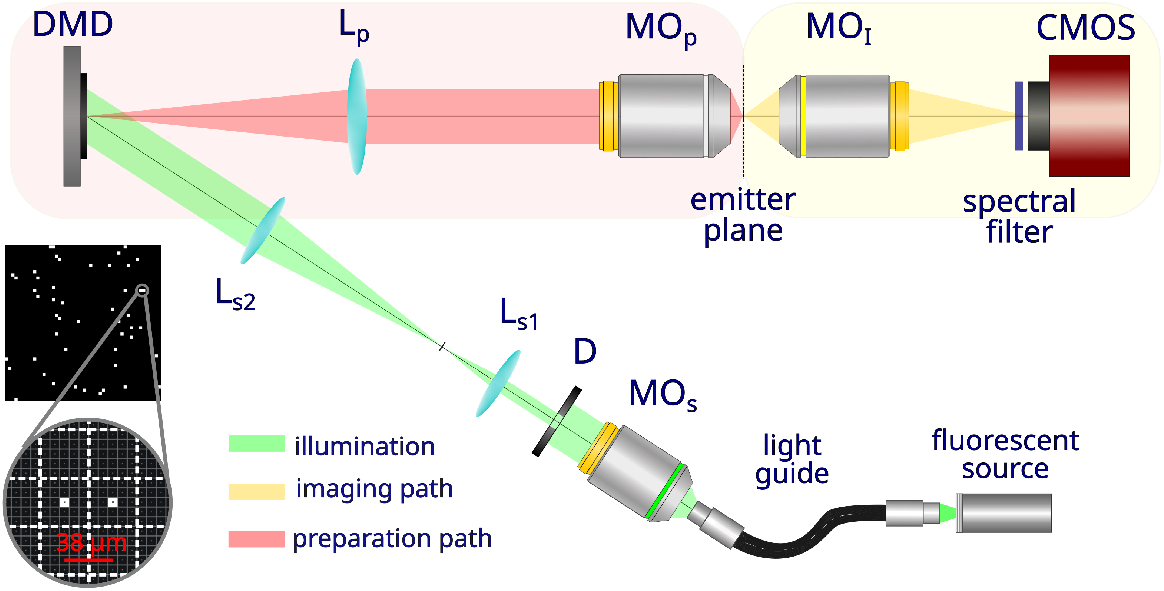
A detailed visualization of the optical setup. In the illumination path, we generate an incoherent light collimated by a microscopic objective MO_s_. The subsequent iris diaphragm D and two achromatic doublets, L_s1_ and L_s2_, improve the illumination homogeneity and reduce undesired reflections from the passive DMD parts. In the preparation path, the DMD-induced ground-truth mask is re-imaged by an achromatic doublet L_p_ and a high-resolution microscope objective MO_p_, creating point-like emitters in the emitter plane. Lastly, this plane is projected by a low-resolution microscope objective MO_I_ onto a camera with a mounted spectral filter.

## Notes

https://github.com/VasinkaD/DAMN

https://zenodo.org/records/14641652

### Competing Interest Statement

The authors have declared no competing interest.

### Summary of Updates

Section "Introduction" was revised to "Main" and "Super-resolution and device-dependence" sections. Figure 7 was relocated to Extended Data.

